# Systematic analysis of the molecular architecture of endocytosis reveals a nanoscale actin nucleation template that drives efficient vesicle formation

**DOI:** 10.1101/217836

**Authors:** Markus Mund, Johannes Albertus van der Beek, Joran Deschamps, Serge Dmitrieff, Jooske Louise Monster, Andrea Picco, François Nédélec, Marko Kaksonen, Jonas Ries

## Abstract

Clathrin-mediated endocytosis is an essential cellular function in all eukaryotes that is driven by a self-assembled macromolecular machine of over 50 different proteins in tens to hundreds of copies. How these proteins are organized to produce endocytic vesicles with high precision and efficiency is not understood. Here, we developed high-throughput superresolution microscopy to reconstruct the nanoscale structural organization of 23 endocytic proteins from over 100,000 endocytic sites in yeast. We found that proteins assemble by radially-ordered recruitment according to function. WASP family proteins form a circular nano-scale template on the membrane to spatially control actin nucleation during vesicle formation. Mathematical modeling of actin polymerization showed that this WASP nano-template creates sufficient force for membrane invagination and substantially increases the efficiency of endocytosis. Such nanoscale pre-patterning of actin nucleation may represent a general design principle for directional force generation in membrane remodeling processes such as during cell migration and division.

## INTRODUCTION

Clathrin-mediated endocytosis (CME) is critical for many biological processes including cell signaling, nutrient uptake and pathogen entry. It is a highly efficient process, which involves the internalization of cargo molecules from the cell surface into small membrane vesicles, and follows a stereotypic temporal order of events: First, a protein coat assembles on the membrane, which is then invaginated to form a membrane vesicle with cargo molecules inside. This vesicle is pinched off the plasma membrane and becomes rapidly uncoated to allow subsequent fusion with endosomal compartments. CME is performed by a complex machinery that comprises more than 50 different proteins and is highly conserved from yeast to humans. Much of our current knowledge of the mechanism of CME comes from research in cultured mammalian cells, and yeast (Boettner et al., 2012; McMahon and Boucrot, 2011; Weinberg and Drubin, 2012). In the budding yeast *Saccharomyces cerevisiae*, large-scale genetic and imaging-based screens have led to the identification of many endocytic proteins, and yielded a near-complete parts list (Munn et al., 1995; Raths et al., 1993; Wendland et al., 1996). In addition, live-cell fluorescence imaging has revealed the order of recruitment and disassembly of components, and categorized endocytic proteins into modules based on their dynamics and function (Kaksonen et al., 2003; 2005). This modular organization is remarkably conserved in metazoans (Boettner et al., 2012; McMahon and Boucrot, 2011; Weinberg and Drubin, 2012).

In yeast, CME can be divided into an initial phase, when early endocytic proteins are recruited to a flat membrane (Kukulski et al., 2012), and a late phase during which invagination occurs. The initial phase is characterized by the recruitment of various endocytic adaptor and coat proteins, including clathrin, and is long and variable in duration (Godlee and Kaksonen, 2013; Kaksonen et al., 2005; Reider and Wendland, 2011; Stimpson et al., 2009). The following late phase is however highly regular across all sites. It begins with the arrival of late coat proteins, which are followed by actin regulatory proteins including WASP and type I myosins (Sun et al., 2006). A short burst of actin polymerization starts membrane invagination, and concomitant with the arrival of amphiphysin proteins, vesicle scission occurs (Kaksonen et al., 2003; Picco et al., 2015).

While the timing of recruitment of individual proteins during endocytosis has been determined in detail, their spatial organization at endocytic sites is largely unknown. This is due to the complexity, dynamics and small size of the endocytic machinery, which is below the resolution of conventional light microscopy. Live-cell fluorescence microscopy could reveal the average positions of endocytic proteins along the invaginated plasma membrane (Berro and Pollard, 2014; Picco et al., 2015), while the shape of the invagination was determined by correlative light and electron microscopy (CLEM) (Idrissi et al., 2008; 2012; Kukulski et al., 2012), but this imaging modality does not offer protein specific contrast. Furthermore, immuno-electron microscopy (EM) reported the approximate location of some endocytic proteins (Idrissi et al., 2008; 2012) in the late stages of endocytosis. However, systematic information about the location of the different endocytic proteins throughout the process is currently lacking, particularly during the initial phase before membrane bending. Thus, the molecular architecture of this complex supramolecular machine, and how it can drive endocytosis so efficiently to reproducibly generate precisely shaped and sized membrane vesicles, remains unknown.

To address this fundamental gap in our knowledge, we developed a high-throughput superresolution microscopy (HT-SRM) and data analysis pipeline to study the nanoscale organization of proteins in the macromolecular endocytic machine over the entire endocytic time line. HT-SRM allowed us to achieve a resolution close to EM while taking advantage of the molecular specificity and large field of view of fluorescence microscopy. By automating both image acquisition and data analysis, we could acquire and process superresolution images of many thousands of cells with minimal user intervention. We used budding yeast to enable systematic fluorescent tagging of endocytic proteins at their genomic loci, thereby ensuring high labeling efficiency while retaining native expression and biological function. This approach allowed us to analyze the structural organization of 23 different proteins of the endocytic machinery sampled throughout the endocytic process at in total over 100,000 endocytic sites in more than 20,000 fixed cells.

We found that assembly of the machinery initiates stochastically from irregular structures. Thereafter, an intricate self-organization emerges where endocytic proteins are radially ordered according to their function. WASP family proteins formed a nanoscale template on the flat membrane to pattern the nucleation of actin filaments. Brownian dynamics simulations of actin polymerization showed that this geometry is essential to induce a scaffold of actin filaments producing sufficient force for membrane invagination and, by centering it provides a mechanism for the high efficiency and robustness of vesicle budding.

## RESULTS

### Experimental pipeline

Here, we used single-molecule localization microscopy (SMLM, also referred to as (f)PALM or STORM (Betzig et al., 2006; Hess et al., 2006; Rust et al., 2006)) to image sites of clathrin-mediated endocytosis in budding yeast strains with single endocytic proteins endogenously tagged with a photoswitchable fluorescent protein. We placed the focal plane on the underside of cells, where endocytic invaginations are oriented perpendicularly to the focal plane (Figure 1A top). Thereby we obtained two-dimensional projections of endocytic structures, which reveal the lateral distribution of proteins at endocytic sites. In these images, the distribution of endocytic proteins appeared as patches, rings, or irregular shapes (Figure S1).

**Figure 1:**
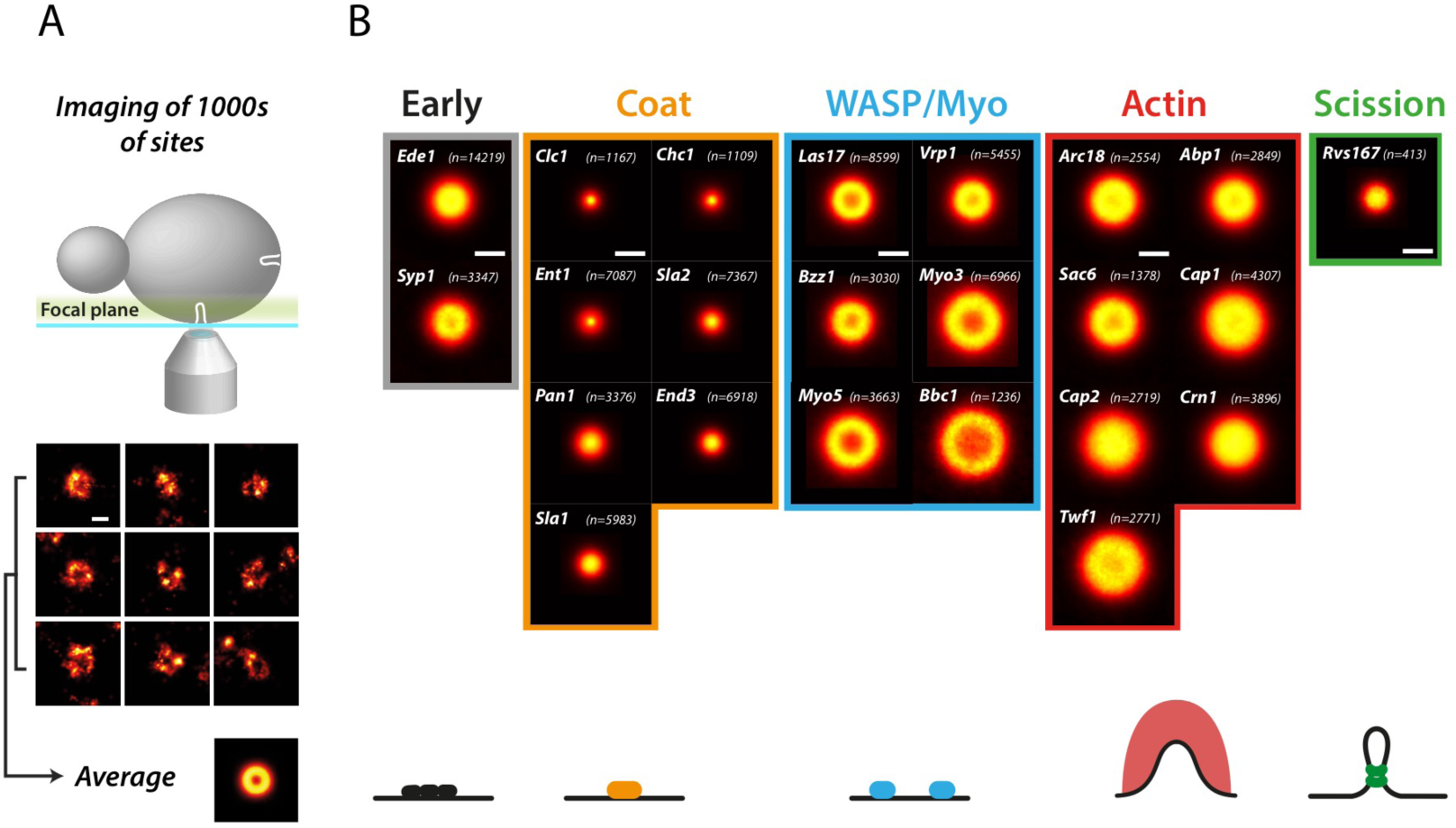
Endocytic proteins are radially organized according to function. A) For each endocytic protein, 1000-10000 cells were imaged using HT-SRM, where the focal plane was placed at the bottom of the cells. Endocytic sites located at the center bottom of the cells were automatically segmented. Shown are nine example sites, and the corresponding average. B) Endocytic proteins form very diverse structures. Shown are the average images of 23 endocytic proteins. The numbers of analyzed endocytic sites are indicated. Scale bar is 100 nm. See also Figure S1.

Because endocytosis was arrested in live cells by rapid fixation, the individual images provide snapshots of different endocytic time points. In order to sample the entire endocytic timeline with high statistical power, we automatically acquired superresolution images of many thousands of endocytic sites, quantitatively analyzed individual structures, and spatially aligned them by translation and subsequently averaged them (Figure 1A bottom). This generated density profiles of how each protein is on average distributed around the center of the endocytic site (Figure 1B), representing the average structural organization of endocytic proteins over their lifetime.

### The functional modules of endocytosis occupy distinct radial zones

Using the imaging pipeline described above, we determined the structural organization of a selection of 23 endocytic proteins from all functional modules of the machinery (Figure 1B, Figure S1). These were early proteins Ede1 and Syp1, coat proteins Sla1, Sla2, Clc1, Chc1, Ent1, End3 and Pan1, WASP/Myo module proteins Las17, Vrp1, Bzz1, Bbc1, Myo3 and Myo5, actin module proteins Abp1, Arc18, Cap1, Cap2, Sac6, Twf1 and Crn1, and the amphiphysin Rvs167. For each of these proteins, we imaged ~1,000-6,000 cells, which yielded superresolution images of ~1,000-10,000 endocytic sites (see table S1). We then automatically derived each protein’s average radial density profile (Figure S1), and classified the shape of the protein distribution as a patch, ring or dome (table S1).

This comprehensive dataset revealed that endocytic proteins assemble into nanoscale structures of strikingly different sizes and geometries, which was evident both from images of individual endocytic sites (Figure 1A, Figure S1) as well as from the average distributions (Figure 1B, Figure S1). Proteins within the same functional endocytic module exhibited similar radial profiles and sizes (Figure 2), indicating that the structural organization of endocytic proteins is predictive of their function. Coat proteins formed the smallest structures, which were mostly patches with a mean outer radius of ~25-50 nm. Proteins of the WASP/Myo module formed larger, ring-like structures with outer radii of ~60-100 nm, and proteins in the actin module formed the largest structures with outer radii of ~80-100 nm. The amphiphysin Rvs167, which is recruited to the invagination around the time of scission, formed small structures with an average outer radius of ~40 nm. Together, our results reveal that endocytic proteins are functionally organized in radial zones at endocytic sites.

**Figure 2:**
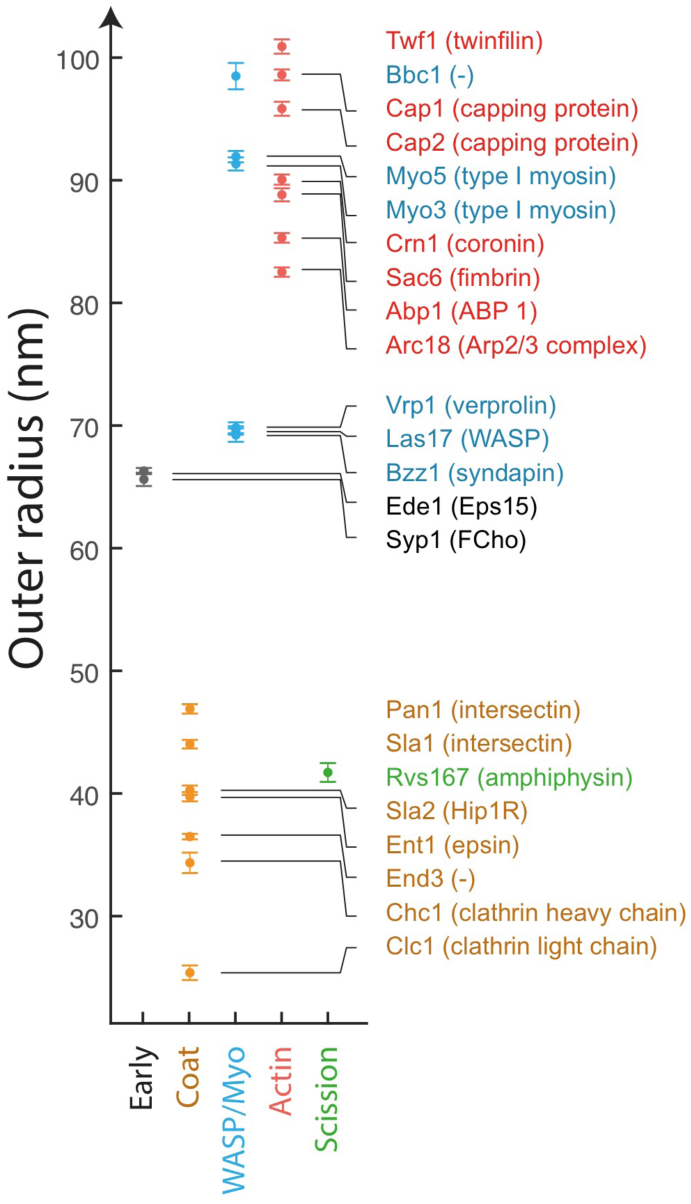
Average outer radii of endocytic proteins. Mean outer radii of structures formed by 23 endocytic proteins from all five functional modules. Error bars indicate SEM. Mammalian homologues are indicated in parentheses, as identified from (Weinberg and Drubin, 2012). Scale bar is 100 nm. See also Figure S1.

### Initiating proteins form irregular structures that grow over time

The individual images of most endocytic proteins resembled their average distribution, suggesting that their organization is fairly constant during endocytosis. However, the earliest proteins, such as Ede1 and Syp1 that initiate endocytic sites with variable timing on the order of minutes and on average form medium-sized structures with outer radii of ~66 nm (Figure 3A), behaved very differently. Ede1 and Syp1 structures showed high variability and included rings, patches, crescents, lines and more irregular shapes of various sizes (Figure S1). To determine whether this heterogeneity corresponded to structural changes over time, we combined superresolution imaging of the early scaffold protein Ede1 with diffraction-limited imaging of a GFP-labelled coat protein Sla2, whose abundance increases over time until vesicle scission (Picco et al., 2015), providing a reference to sort endocytic sites in time (Figure 3B). We found that the distribution of Ede1 grows in size over time, with an increase in median outer radius from 57.5 ± 0.4 nm (mean ± standard error of the mean) up to 67.9 ± 0.5 nm (Figure 3C-E). These data indicate that the self-organizing endocytic machinery is initiated stochastically with variable structures, which are remodeled during coat recruitment.

**Figure 3:**
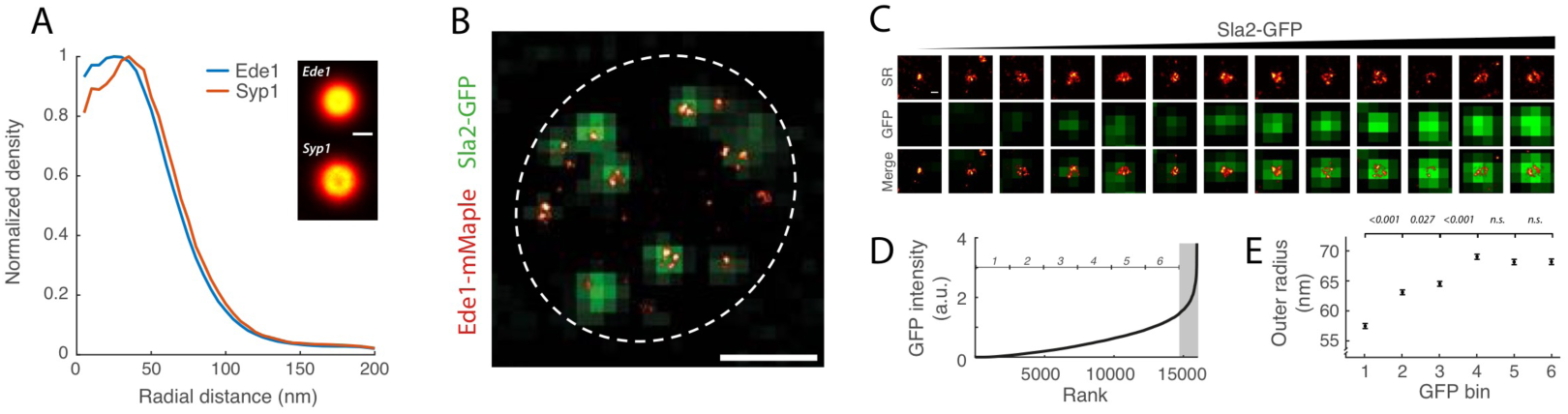
Early proteins form diverse structures that grow over time. A) Average radial distribution of Ede1 and Syp1. B) Staging of endocytosis by combining superresolution imaging of Ede1 with diffraction-limited imaging of Sla2-GFP. C) Example sites of Ede1 for increasing Sla2-GFP intensity. D) Quantification of GFP intensity at Ede1 sites. Endocytic sites were sorted by ascending Sla2-GFP signal, and binned in six time windows. The shaded area indicates a high fraction of sites with overlapping GFP signals, e.g. in small buds, which were thus excluded from the analysis. E) Outer radii (mean ± SEM) of Ede1 over time. P values are indicated. Scale bars are 100 nm (A, C) and 1 µm (B).

### Coat proteins assemble from center to periphery in growing patches

Besides Sla2, a variety of other coat proteins are recruited to the endocytic site. Here, we determined the average distributions of the coat proteins Clc1, Chc1, Ent1, Sla1, Sla2, Pan1 and End3, and found that they formed patch-like structures with sizes that were significantly different between each other (p < 0.0005 for all pairwise size differences) in the range between 25-50 nm (Figure 4A). Clathrin (Clc1 and Chc1), which arrives first, formed the smallest structures, followed by End3, Ent1 and Sla2. The late coat proteins Sla1 and Pan1 formed the largest structures. Although the overall averages of all coat proteins were patch-like, a fraction of the individual sites of the late proteins showed ring-like distributions (Figure S1). This is in agreement with our previous findings that Sla1 can form small rings at endocytic sites (Picco et al., 2015). Thus, HT-SRM identified clear organizational differences between endocytic coat proteins, suggesting that their progressive assembly translates into a radially expanding architecture around the endocytic sites: the early arriving clathrin, Sla2 and Ent1 form at the core of the coat while the late coat proteins Sla1 and Pan1 extend to its periphery.

**Figure 4:**
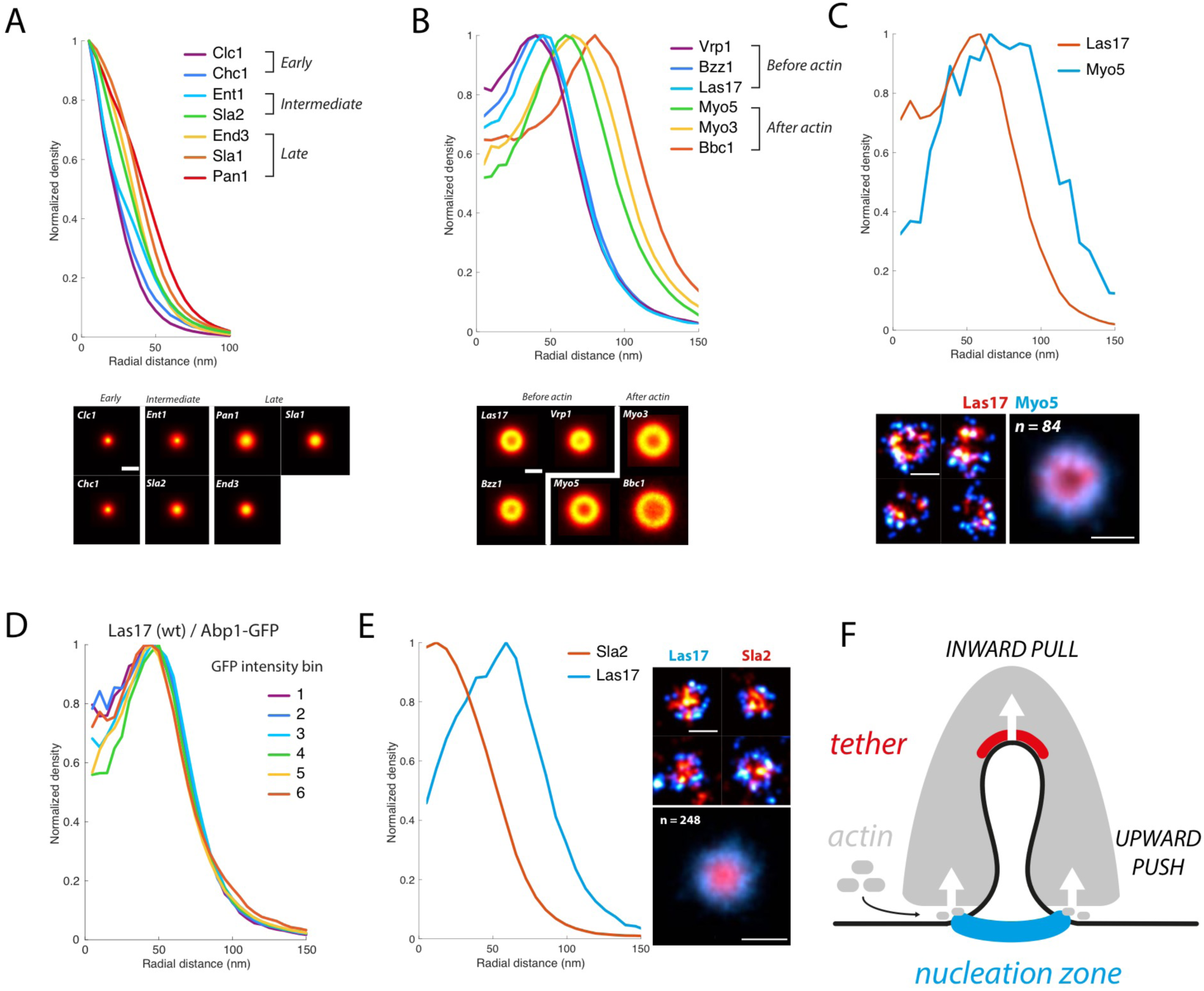
Actin nucleation is pre-patterned by WASP around the coat. A) Average images and radial profiles of coat proteins. B) Average images and radial profiles of WASP/Myo proteins. C) Dual-color superresolution images of Las17 and Myo5 at individual endocytic sites, and the average radial distribution derived from 84 sites. D) Average radial profiles of Las17 over time, as determined using Abp1-GFP as a diffraction-limited timing marker. E) Dual-color superresolution images of Las17 and Sla2 at individual endocytic sites, and the average radial distribution derived from 284 sites. F) Model for force generation by the actin network. The addition of actin monomers at the bottom pushes up the entire network, which is tethered to the plasma membrane in the center and thus induces its invagination. Scale bars are 100 nm. See also Figures S2, S3, S4.

### Nucleation promoting factors and their inhibitors form a circular nano-template for actin polymerization

As the coat is assembling, members of the WASP/Myo module, which regulate actin nucleation, start to arrive at endocytic sites. The average radial profiles of these actin-regulatory proteins revealed that they formed rings of different sizes (Figure 4B). The group of Las17 (yeast WASP), Bzz1 and Vrp1 formed rings with indistinguishable radii of ~70 nm, in agreement with their reported interactions and functional cooperation to promote actin nucleation (Evangelista et al., 2000; Grötsch et al., 2010; Kishimoto et al., 2011; Lewellyn et al., 2015; Sun et al., 2006). By contrast, the rings of the type-I myosins Myo3 and Myo5 were significantly larger with a radius of 90 nm. This difference was confirmed in dual-color superresolution images of individual endocytic sites, where Myo5 was peripherally arranged around a Las17 ring (Figure 4C). The fact that the Myo3/5 radii were significantly larger than Vrp1, which is necessary to recruit them (Grötsch et al., 2010; Lewellyn et al., 2015) may indicate that the myosins first bind to actin and only subsequently interact with Vrp1.

The WASP protein Las17 is known to arrive at endocytic sites together with the most peripheral coat protein Pan1 around 20 seconds before the onset of actin polymerization. Strikingly, Las17 is therefore the first protein to form clear ring-like structures. We therefore asked if Las17 is already pre-patterned as a ring on the flat plasma membrane, or whether its ring-like distribution is caused by the already invaginating membrane. To test this, we first inhibited actin polymerization with Latrunculin A, which arrests endocytosis prior to membrane bending (Kukulski et al., 2012). Under these conditions, Las17 still formed rings of slightly larger size (outer radius of 81.6 ± 30 nm, Figure S2), demonstrating that ring structures of nucleation promoting factors can form on the flat membrane in the absence of polymerized actin. Next, we analyzed whether Las17 rings changed in size during invagination by imaging Las17 in super-resolution alongside Abp1-GFP as a temporal reference for actin polymerization (Figure S3). We found that the radii of Las17 rings as well as the fraction of ring-like structures were constant over time (Figure 4D). We conclude that Las17 forms a robust ring structure with a radius of ~70 nm early on the flat plasma membrane before actin polymerization begins.

Why then does Las17 form a ring, when the coat proteins assembling before it are forming patches, and actin polymerization has not started to invaginate the membrane? The core and mid layer coat proteins, clathrin, Sla2 and Ent1 are known to form a tight molecular lattice *in vitro* (Skruzny et al., 2015), and could thus potentially prevent the access of later arriving proteins to the center of the endocytic site and determine the minimal size of the ring they form. Consistent with this hypothesis, dual-color superresolution imaging of Las17 and the coat protein Sla2 indeed showed that the average radial density of Sla2 was located precisely inside the Las17 rings (Figure 4E, Figure S4). Thus, our data revealed that actin nucleation is pre-patterned, already on a flat membrane, by a ring of Las17 molecules around the center of the endocytic site where membrane-actin coupling is provided by Sla2 and Ent1/2. Actin monomers are thus added to filaments close to this nucleation zone. This induces an upward push of the actin network, which is attached to the membrane in the center, explaining how polymerizing actin can pull the plasma membrane inwards (Figure 4F).

Interestingly, coat proteins may not only set the physical inner limit of actin nucleation by Las17, but may also ensure this boundary biochemically, as the peripheral coat protein Sla1 is a WASP inhibitor (Rodal et al., 2003). This prompted us to ask if there is also an outer limit of actin nucleation. Bbc1, another WASP inhibitor (Rodal et al., 2003), formed the largest rings of the WASP/Myo module with a radius of ~98 nm. Thus, the ring of Las17 is sandwiched between an inner patch of its inhibitor Sla1 and an outer ring of its inhibitor Bbc1, which could precisely limit Las17 activity for actin nucleation to a narrow radial zone at around 70 nm (Figure 5A).

**Figure 5:**
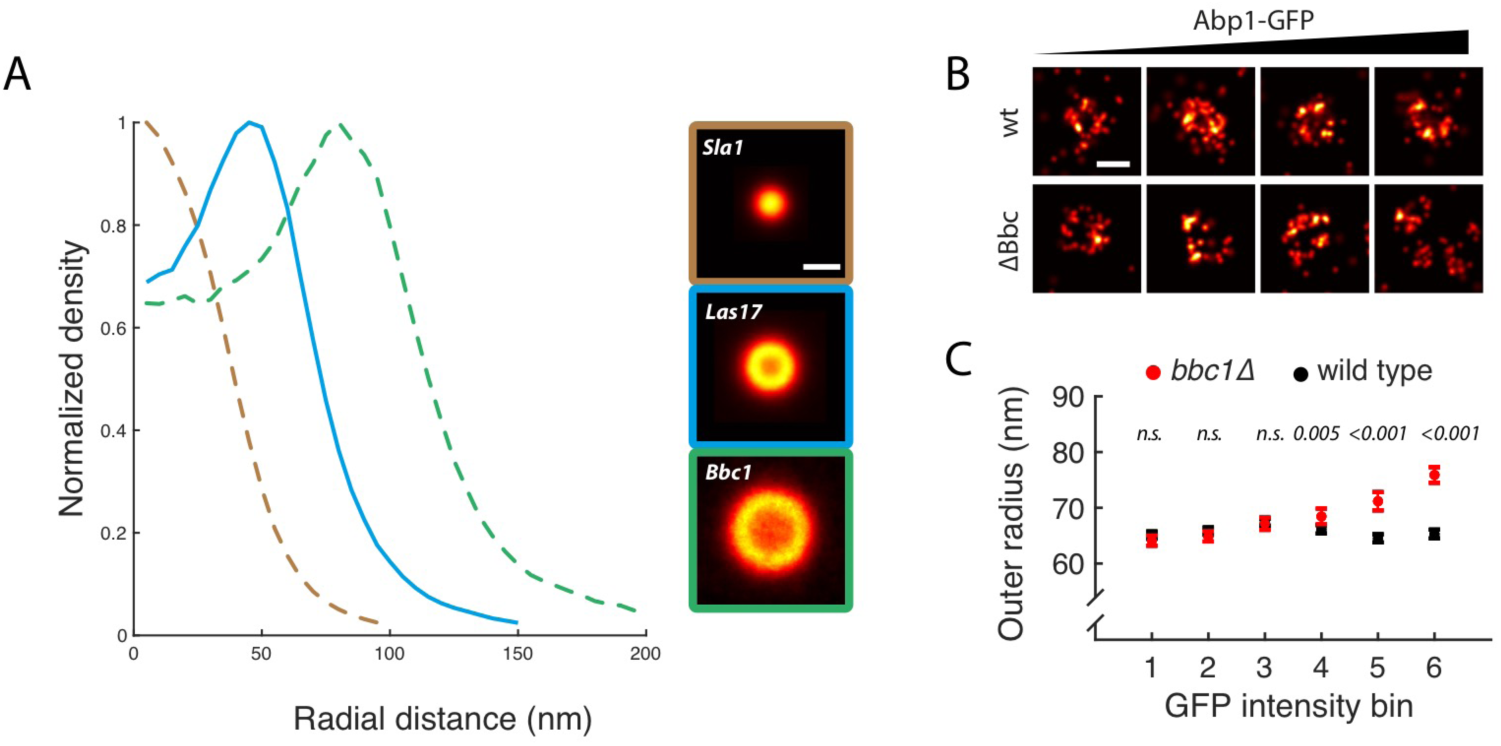
WASP is sandwiched by its inhibitors. A) Radial profiles of Las17 and its inhibitors Sla1 and Bbc1. B) Individual Las17 sites in wild type and Bbc1 deletion backgrounds, with the corresponding Abp1-GFP signal increasing from left to right. C) Outer radii of Las17 in wild type and Bbc1 deletion backgrounds for increasing Abp1-GFP intensity. p values from pair-wise t-tests between wild type and Bbc1 deletion for the respective time points are indicated. Scale bar is 100 nm.

To test if this narrowly sandwiched zone of Las17 localization and activity is functionally important, we perturbed its structure. To this end, we deleted Bbc1 and analyzed the distribution of Las17 over time using the actin marker Abp1 as a temporal reference for actin polymerization. When Bbc1 was deleted, Las17 initially formed the same sized rings as in the wild type. However once actin polymerization began, Las17 structures expanded significantly (mean outer radius ± SEM from 64.2 ± 0.9 to 76.0 ± 1.4 nm, p < 10^−6^) and eventually fragmented (Figure 5B, C). Thus, upon actin polymerization at the endocytic site, Las17 localization is indeed regulated by Bbc1.

Taken together, we have discovered that the major WASP family actin nucleation promoting factor Las17 is sandwiched by its inhibitors on the inside and outside to form a nanotemplate for actin nucleation from a ring of about 70 nm in radius. This precise molecular architecture is established on the flat plasma membrane well before the onset of actin polymerization. During actin polymerization the Las17 ring is stabilized by Bbc1.

### Temporal reconstruction shows dynamic nucleation of actin from the WASP nucleation zone

During the late phase of endocytosis, the plasma membrane is invaginated by the formation of an actin network. Here, we first determined the average radial distribution of the seven actin network components Abp1, Arc18, Cap1, Cap2, Crn1, Sac6 and Twf1. We found that they all formed similar large structures with narrow intensity minima in the center (Figure 6A), matching their expected distribution in a dome-like actin network that encompasses the endocytic membrane invagination (Idrissi et al., 2008; Kukulski et al., 2012; Mulholland et al., 1994). Although most of these proteins are recruited at the same time with highly similar dynamics (Kaksonen et al., 2005; Picco et al., 2015), their nanoscale radial profiles revealed differences in their outer radii, indicating that the barbed ends of actin filaments protrude outwards further than pointed ends (Figure S5).

**Figure 6:**
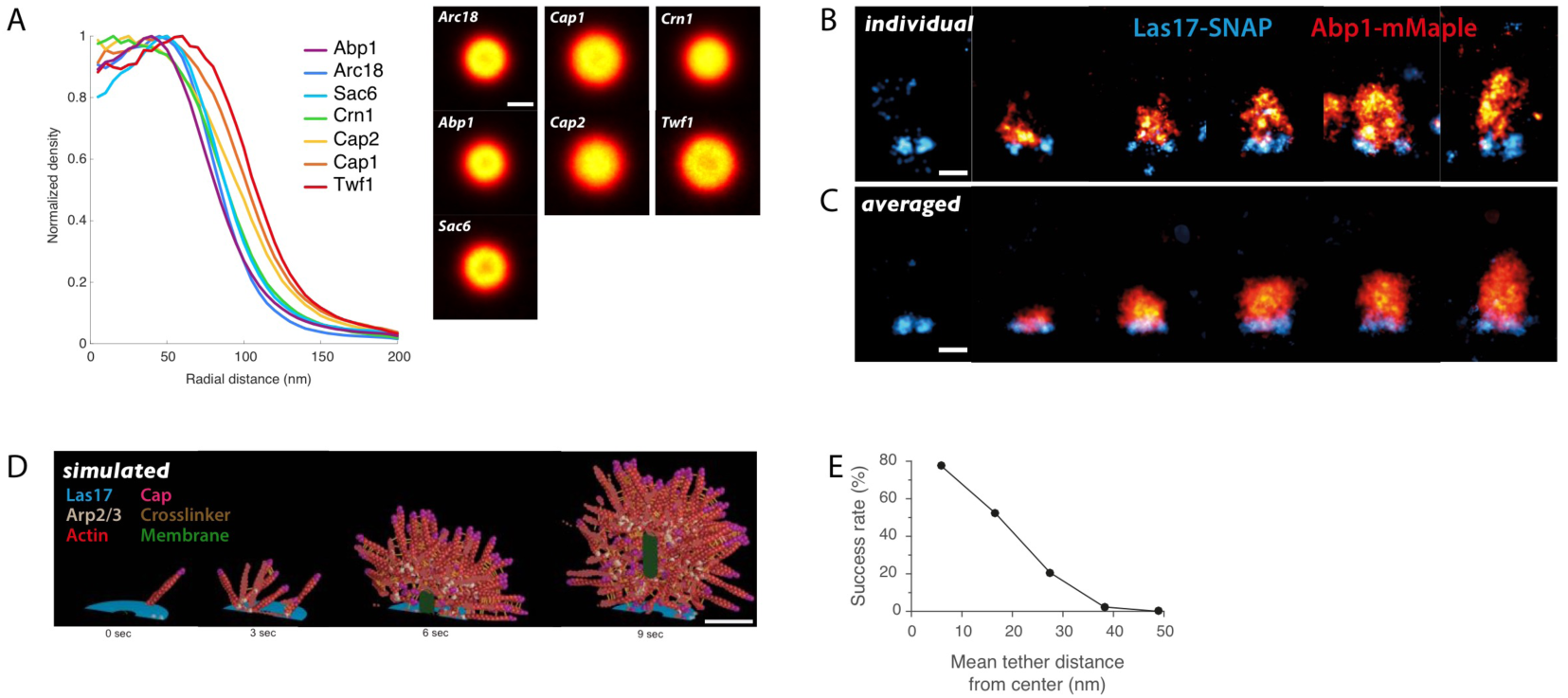
The actin network emanates from the WASP nucleation zone. A) Average images and radial profiles of proteins from the actin module. B) Selection of dual-color superresolution images of Las17 and Abp1 at individual sites from a side-view perspective. Images have been rotated so that the direction of endocytosis is upwards, and sorted by the distance of Abp1 centroid to Las17 at the base. C) Running-window averages of Las17 and Abp1 at endocytic sites. D) Simulations of the actin network at endocytic sites using Cytosim. The invagination was described as a solid vesicle progressing inwards when pulled by actin, which is connected to the top of the vesicle, mimicking the activity of Sla2 and Ent1/2 proteins. New actin filaments were nucleated by Las17 stimulated Arp2/3 complex, grew at 1 µm/sec with a stall force of ~10 pN, and were capped at a length of 60 nm (for details see methods). Shown are four intermediate states. Only half of the network is displayed for clarity. E) Plot of endocytic success rate as a function of lateral position of the invagination with respect to the nucleation zone. Scale bars are 100 nm. See also Figures S5, S6, S7.

Next, we wanted to directly visualize the spatial relationship of the Las17 nucleation zone and the resulting actin network. To this end, we used dual-color superresolution microscopy of Las17 and the actin marker Abp1 in a side-view configuration (Figure 6B). While Las17 always localized close to the base of the plasma membrane, Abp1 formed structures of very different shapes and sizes, consistent with the dynamic nature of actin polymerization during endocytosis. To infer the endocytic time of our snapshots, we used previously reported live-cell data that the center of mass of Abp1 is continuously moving inwards during endocytosis (Figure S6A, (Picco et al., 2015)). Thus, by sorting our snapshots by increasing distance between the centroids of the Las17 and Abp1, we sorted them in time. The resulting temporal reconstruction (raw data or running window averaging) revealed that the actin network emanates directly above Las17 and progressively grows into the cytoplasm (Figure 6B,C). The average structures showed good agreement with the previously reported dimensions of the actin network (ranging from ~100 nm to 200 nm in width, and 70 nm to 240 nm in height, for early and late time points respectively) as estimated using CLEM (Kukulski et al., 2012) (Figure S6B), validating our approach. In summary, ordering our snapshot data in time directly shows how the Las17 nanotemplate guides actin polymerization.

### Brownian dynamics simulations show that symmetric actin polymerization around the membrane invagination increases the efficiency of endocytosis

During endocytosis, actin assembly overcomes the turgor pressure, which opposes membrane invagination with forces exceeding 1000 pN (Basu et al., 2014; Dmitrieff and Nédélec, 2015), much higher than the polymerization stall force of single actin filaments of 1-10 pN (Dmitrieff and Nédélec, 2016). Nevertheless vesicles form with very high efficiency and uniform size from endocytic patches (Kaksonen et al., 2003; Kukulski et al., 2012). We thus speculated that the nanoscale patterning of actin nucleation that we discovered could be important to allow F-actin to generate the force required to form a vesicle. To test this, we simulated the mechanics of the dynamic endocytic actin network using the open-source modeling framework Cytosim (Nédélec and Foethke, 2007). We used 250 growing filaments based on previously measured molecular numbers of Arp2/3 (Picco et al., 2015) and used previously reported, realistic parameters for membrane bending forces (Dmitrieff and Nédélec, 2015). We then simulated a minimal set of components including Arp2/3, passive crosslinkers, and Las17 patterned as observed. Within these simulations, actin filaments self-organized into a 3D branched network of realistic dimensions and molecular composition that could reliably produce forces exceeding 1000 pN, and thus was able to overcome turgor pressure (Figure 6D, Movie S1). The methods section includes a detailed description of the model.

This computational model allowed us to tested whether a ring-shaped nucleation zone is advantageous for the efficiency of vesicle budding, as many different geometries might produce a sufficiently strong actin network capable of overcoming turgor pressure. A key advantage of a ring-shaped nucleation zone could be that membrane invagination occurs in its center, with actin polymerizing symmetrically around it, preventing lateral displacement or tilting of the invagination. Indeed, our simulations showed that ensuring actin nucleation occurs all around the invagination, rather than allowing actin filaments to be nucleated asymmetrically with respect to the invagination, had a strong effect on vesicle budding efficiency. Specifically, if actin filaments were nucleated around a central invagination zone of 25 nm radius consistent with the hole we observed in Las17 rings, the membrane invagination remained central with a mean position of below 11 nm away from the center and endocytosis had a 80% chance of success. If the actin nucleation zone was positioned asymmetrically, the success rate of endocytosis dropped substantially, and endocytosis was entirely unsuccessful when the center of nucleation activity was on average more than 43 nm away from the invagination (Figure 6E). These results indicate that actin-driven pulling on the membrane is substantially more efficient in creating invaginations into the cytoplasm if they are confined to a small area in the center of a ring. In addition, off-centered invaginations often moved sideways, rather than being elongated, with the actin forming distinct comets (Figure S7, Movie S2), which is not observed in vivo. We conclude that the ring-like nanotemplate of actin nucleation ensures high efficiency of vesicle budding by confining the membrane invagination at its center and promoting actin polymerization symmetrically around it, which directs vesicle movement perpendicular to the plasma membrane.

### Discussion

Here, we have established an automated HT-SRM pipeline to record and quantitatively analyze superresolution images of thousands of yeast cells without user intervention. The capability to image large numbers of cells brought multiple advantages: it lent strong statistical power to the study, allowed fine sampling of the structural space (as seen for Ede1, which first assembles into irregular structures), and the detection of rare events (as seen for Rvs167, which has a very short lifetime). Thus, automation enabled us to extend the scope of this study to image 23 endocytic proteins, and to capture the structural variability of the highly dynamic endocytic machinery by imaging more than 100,000 endocytic sites.

### Spatial organization correlates with function and reveals a radial assembly of the machinery

We discovered that the endocytic machinery has an intricate radial organization, where proteins occupy distinct radial zones. Early proteins first form diverse structures, which become bigger and more regular over time. Coat proteins form a patch structure in the center of the endocytic site, around which WASP/Myo proteins populate ring-shaped zones of three distinct sizes. Actin-binding proteins occupy the largest radial area, reflecting that the actin network forms a dome enveloping the invagination. Around the time-point of scission, amphiphysins form small structures compatible with binding to the narrow neck of the invagination.

Generally, radial dimensions increased over the endocytic time line. Given that proteins from the early, coat and WASP/Myo modules are recruited directly or indirectly to the flat plasma membrane prior to actin polymerization, our data suggest that newly arriving proteins bind to already recruited components at the periphery in the plane of the membrane, revealing the mechanism by which the endocytic machinery is collectively assembled (Figure 7).

**Figure 7:**
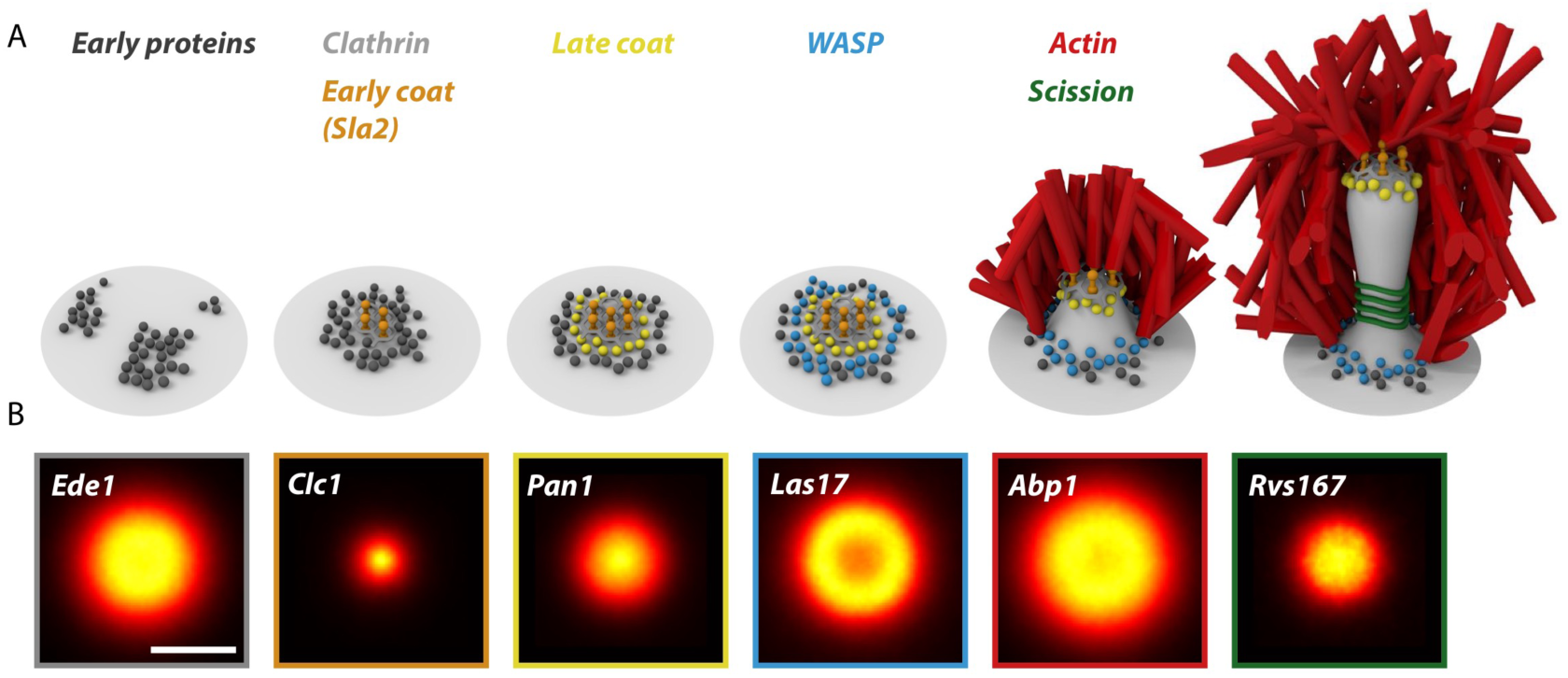
The endocytic machinery assembles by peripheral binding. A) Schematic representation of the assembly of the endocytic machinery. B) Representative radial averages of endocytic proteins of key steps during assembly. Shown are Ede1 (early), Clc1 (Coat early), Pan1 (Coat late), Las17 (WASP), Abp1 (Actin), Rvs167 (Scission). Scale bar is 100 nm.

The coat occupies the center of the endocytic site. Sla2 and Ent1/2 bind cooperatively to the membrane, and to clathrin, forming a lattice (Skruzny et al., 2015). We showed that the subsequently arriving late coat proteins Pan1 and Sla1 extend the patch in size. This structure is likely stable, which is corroborated by slow molecular exchange of these proteins, at least when endocytosis is arrested by LatA treatment (Skruzny et al., 2012). Taken together, these results support an assembly model whereby the compactness of the coat inhibits penetration by subsequently recruited proteins, which thus bind to its periphery.

Coat assembly is itself preceded by the arrival of the early proteins Ede1 and Syp1, which our study reveals to form a variety of irregular shapes and well-defined rings. The radial profiles of Syp1 and Ede1 were similar, which is expected as they directly interact already in the cytoplasm, and Syp1 recruitment requires Ede1 (Boeke et al., 2014; Stimpson et al., 2009). We showed that Ede1 assemblies increase in size over time, which agrees with increasing molecular numbers of Ede1 (Stimpson et al., 2009). The formation of bigger, more ring-like structures around the coat may be induced by the competition of Ede1 with late coat proteins for binding to the early coat proteins Ent1/2 via EH domains both in Ede1 and Pan1 (Boeke et al., 2014; Wendland and Emr, 1998).

We found that after Sla2 recruitment commences, Ede1 structures became rather constant in size. We thus propose that the beginning of coat recruitment represents a structural switch of the endocytic machinery, which correlates with the transition into the late phase of endocytosis that leads to vesicle budding with highly stereotypic timing (Kaksonen et al., 2005).

Within the WASP/Myo module, recruitment time and ring size were again correlated: a first set of proteins comprising Las17, Vrp1 and Bzz1, all of which directly interact (Sun et al., 2006), formed rings around the coat. Later, concomitant with the onset of actin polymerization, Myo3/5 formed larger, and Bbc1 the largest rings around. Taken together, the increasing ring sizes suggest that over time, endocytic proteins are recruited to distinct (radial) zones, which are determined by a variety of interactions with proteins that are already present at the endocytic site (Figure 7). In the long, variable phase of site initiation, protein clustering due to multiple interactions between proteins and the plasma membrane eventually triggers the formation of a stable coat structure, and endocytosis transitions into a more regular phase.

### A nano-pattern of actin nucleation explains efficient and robust membrane invagination

In yeast, actin polymerization provides the necessary force to bend the plasma membrane, which is particularly high compared to mammalian cells because of its intracellular turgor pressure (Aghamohammadzadeh and Ayscough, 2009; Basu et al., 2014). We showed that actin nucleation, which is rate-limiting for actin polymerization (Pollard and Cooper, 2009), is patterned by Las17, which early on forms rings at the base of the plasma membrane. Using a dynamic reconstruction, we directly visualized how an actin network is formed from this nucleation nano-template.

What are the mechanistic implications of a ring-shaped nucleation zone, and of its establishment early on during endocytosis? We simulated actin polymerization at endocytic sites using Cytosim, and found that centering the nucleation zone around the invagination is crucial for endocytic success. When actin polymerization is patterned asymmetrically around the membrane tethers, comet-shaped actin networks are formed, which are reminiscent of mutant phenotypes when the actin-membrane tether Sla2 is deleted (Kaksonen et al., 2003), and induce lateral displacement rather than membrane invagination.

In light of the inherent stochasticity of macromolecular machineries, the robustness and efficiency of endocytosis in yeast is remarkable. Once actin polymerization begins, endocytosis is virtually always productive (Kaksonen et al., 2005) and occurs with high spatio-temporal regularity and directionality (Berro and Pollard, 2014; Picco et al., 2015). Interestingly, during the late phase of endocytosis, high forces are required already shortly after membrane bending has begun, when the invaginations are still shallow (Dmitrieff and Nédélec, 2015; Hassinger et al., 2017; Walani et al., 2015). Beyond this peak, the force requirement drops substantially and remains comparably low. This corresponds to a snap-through transition (Walani et al., 2015), which could explain the high efficiency of endocytosis. To provide this maximum force already at the start of endocytosis, it seems crucial to optimize the ability to rapidly nucleate actin filaments once this process is initiated. Our data show that actin nucleation is pre-patterned around the center of the endocytic site, where membrane-actin coupling is provided by Sla2 and Ent1/2 (Figure 4F). This provides strong nucleation capacity from the start, centers the membrane tethers, and lends directional stability to the actin machinery. Together, these factors contribute to the regularity of yeast endocytosis during its most crucial phase corresponding to the beginning of invagination.

How do changes in size and shape of the nucleation zone influence geometry and properties of the resulting actin network? When we deleted Bbc1, Las17 initially formed rings as in wild type cells, but with the onset of actin polymerization became larger and less regular. This suggests that Bbc1 not only inhibits the biochemical activity of Las17, which has been reported *in vitro* (Rodal et al., 2003), but also that Bbc1 regulates the spatial distribution of Las17. Interestingly, it was previously shown that in a Bbc1 deletion background, the centroids of Sla1 and Abp1 move inwards further than in wild type cells, and exhibit a leaping movement (Kaksonen et al., 2005). A larger Las17 nucleation zone provides an explanation for this phenotype, as a larger actin network can propel the forming vesicle further into the cytoplasm.

Our discovery of nano-patterned actin filament nucleation is likely to be of general importance in many other systems. These include mammalian cells, where actin rings at the base of the plasma membrane could also be the consequence of the nucleation being patterned (Li et al., 2015). More generally, localized actin nucleation by various nucleation promoting factors is found in many cellular contexts that depend on polarized actin networks of various architectures, including lamellipodia of migratory cells, filopodial membrane extensions, various sites during cell division, and the surface of pathogens, which use actin to propel themselves through the host cell (Campellone and Welch, 2010; Firat-Karalar and Welch, 2011; Pollard and Borisy, 2003; Pollard and Cooper, 2009; Skau and Waterman, 2015). These examples illustrate a variety of nucleation geometries, and show that spatially controlling nucleation is a common strategy to form actin networks that can build up directional forces. It is tempting to speculate that also in these contexts, nucleation is precisely patterned on the nanoscale.

Finally, we hope that our superresolution dataset will be useful to our community, forming a solid foundation for further mechanistic modelling of the functioning of the endocytic machinery.

## AUTHOR CONTRIBUTIONS

M.M., M.K. and J.R. conceived the project and designed the research. M.M., J.A.B., J.L.M. and J.R. conducted experiments and analyzed data. M.M., J.D. and J.R. developed the HT-SRM microscope and online analysis software. S.D. and F.N. designed and performed simulations. A.P. discussed research design, results and the analysis pipeline. M.M. and J.R. wrote the manuscript with input from all authors.

## ACKNOWLEDGEMENTS

We thank Jan Ellenberg for critically reading the manuscript. We thank the entire Ries and Kaksonen labs for fruitful discussions and support of this work. We thank Rohit Prakash and Sunil Dogga for their support during the early phase of the project. We thank all contributors to Cytosim and EMBL IT support team, particularly those in charge of high performance computing. We thank Life Science Editors for editing assistance, and Effigos AG for 3D graphics.

This work was funded by the European Molecular Biology Laboratory (M.M., J.A.vdB., J.M., J.D., S.D., F.N., J.R.), the EMBL International PhD Programme (M.M., J.D.), the EMBL Interdisciplinary Postdoctoral Programme (S.D.), the Deutsche Forschungs Gemeinschaft (DFG RI 2380/2; J.D., J.R.), the Swiss National Science Foundation (31003A_163267; MK) and the European Reseach Council (ERC CoG-724489 — CellStructure; M.M., J.R.).

